# Generative network models identify biological mechanisms of altered structural brain connectivity in schizophrenia

**DOI:** 10.1101/604322

**Authors:** Xiaolong Zhang, Urs Braun, Anais Harneit, Zhenxiang Zang, Lena S. Geiger, Richard F. Betzel, Junfang Chen, Janina Schweiger, Kristina Schwarz, Jonathan Rochus Reinwald, Stefan Fritze, Stephanie Witt, Marcella Rietschel, Markus M. Nöthen, Franziska Degenhardt, Emanuel Schwarz, Dusan Hirjak, Andreas Meyer-Lindenberg, Danielle S. Bassett, Heike Tost

## Abstract

**Background:** Alterations in the structural connectome of schizophrenia patients have been widely characterized, but the mechanisms leading to those alterations remain largely unknown. Generative network models have recently been introduced as a tool to test the biological underpinnings of the formation of altered structural brain networks.

**Methods:** We evaluated different generative network models to investigate the formation of structural brain networks in healthy controls (n=152), schizophrenia patients (n=66) and their unaffected first-degree relatives (n=32), and we identified spatial and topological factors contributing to network formation. We further investigated the association of these factors to cognition and to polygenic risk for schizophrenia.

**Results:** Structural brain networks can be best accounted for by a two-factor model combining spatial constraints and topological neighborhood structure. The same wiring model explained brain network formation for all groups analyzed. However, relatives and schizophrenia patients exhibited significantly lower spatial constraints and lower topological facilitation compared to healthy controls. The model parameter for spatial constraint was correlated with the polygenic risk for schizophrenia and predicted reduced cognitive performance.

**Conclusions:** Our results identify spatial constraints and local topological structure as two interrelated mechanisms contributing to normal brain development as well as altered connectomes in schizophrenia. Spatial constraints were linked to the genetic risk for schizophrenia and general cognitive functioning, thereby providing insights into their biological basis and behavioral relevance.

## INTRODUCTION

Schizophrenia is a highly heritable neurodevelopmental disorder (1–3) characterized by abnormalities in perception, cognition, affect, behavior and social functioning (4). Converging evidence supports the notion that wiring disruptions of brain networks may partially underlie these abnormalities (5–7). Indeed, previous studies have found marked differences in the brain network architecture in schizophrenia (8, 9) and delineated alterations in their structural development (10). More generally, human brain networks show a complex architecture favoring topologically advantageous properties while still being sparsely connected, in line with the proposal that the developmental architecture of the human brain connectome results from an economic trade-off between minimizing wiring costs and allowing adaptively valuable topological features (11). Indeed, there is evidence for alterations in both connection distance (12, 13) and network topology including reduced local clustering and modularity (14, 15) in schizophrenia, consistent with a biased trade-off between wiring cost and topology (11). However, the potential processes by which brain networks develop and their disturbances in schizophrenia are poorly understood.

Current network neuroscience predominantly approaches focus on *descriptive* individual or population-level differences, offering little insight into the mechanisms that give rise to network alterations in brain disorders (16–18). A recent adaption of these methods from the branch of generative network models (GNMs) may help address some of these limitations. GNM-based approaches model the stepwise formation of networks based on wiring rules reflecting potential neurodevelopmental constraints. One can then compare synthetic networks generated by the model to empirical brain networks reconstructed from neuroimaging data, thereby explicitly testing different mechanistic explanations that might govern their (disordered) structural formation (16, 19). Two recent examples have tested different wiring rules in the formation of healthy brain networks, finding converging evidence for a two factor model where one factor accounts for the spatial embedding of brain networks by penalizing spatially distant connections while the other factor enhances a complex local topological organization (20, 21). The model parameters have a high degree of biological plausibility as they account for the metabolic cost of wiring and the strength of a „Hebbian-like“ wiring rule, in part buttressed by the fact that they undergo progressive changes over the lifespan and show alterations in disease states (20, 21).

In addition to their biological plausibility and developmental sensitivity, an appropriate model of brain network architecture might help to illuminate genetic aspects underlying network alterations and formation in mental disorders. An important strategy is the examination of unaffected first-degree relatives of patients, who have an increased familial risk for developing the disorder (22–24). This allows for the identification of intermediate brain phenotypes linked to psychiatric risk independent of potential disease-related confounders (25). In addition, the genetic contributions to these phenotypes can be studied with modern genetic approaches utilizing the potential of cumulative genetic risk scores.

Here, we combined GNMs with imaging genetics analyses to identify potential developmental mechanisms promoting the altered formation of structural brain networks in schizophrenia (Figure 1). Building on a family of previously described and validated generative models (20), we first replicated their best-fitting model in a healthy sample, and we subsequently applied it to a group of unaffected first-degree relatives and schizophrenia patients. Following the hypothesis that there exists an aberrant balance between wiring cost and topological properties during network formation in schizophrenia, we tested whether these model parameters show the quality of an intermediate phenotype, relate to schizophrenia polygenic risk and are relevant for cognitive function.

**Figure 1.**
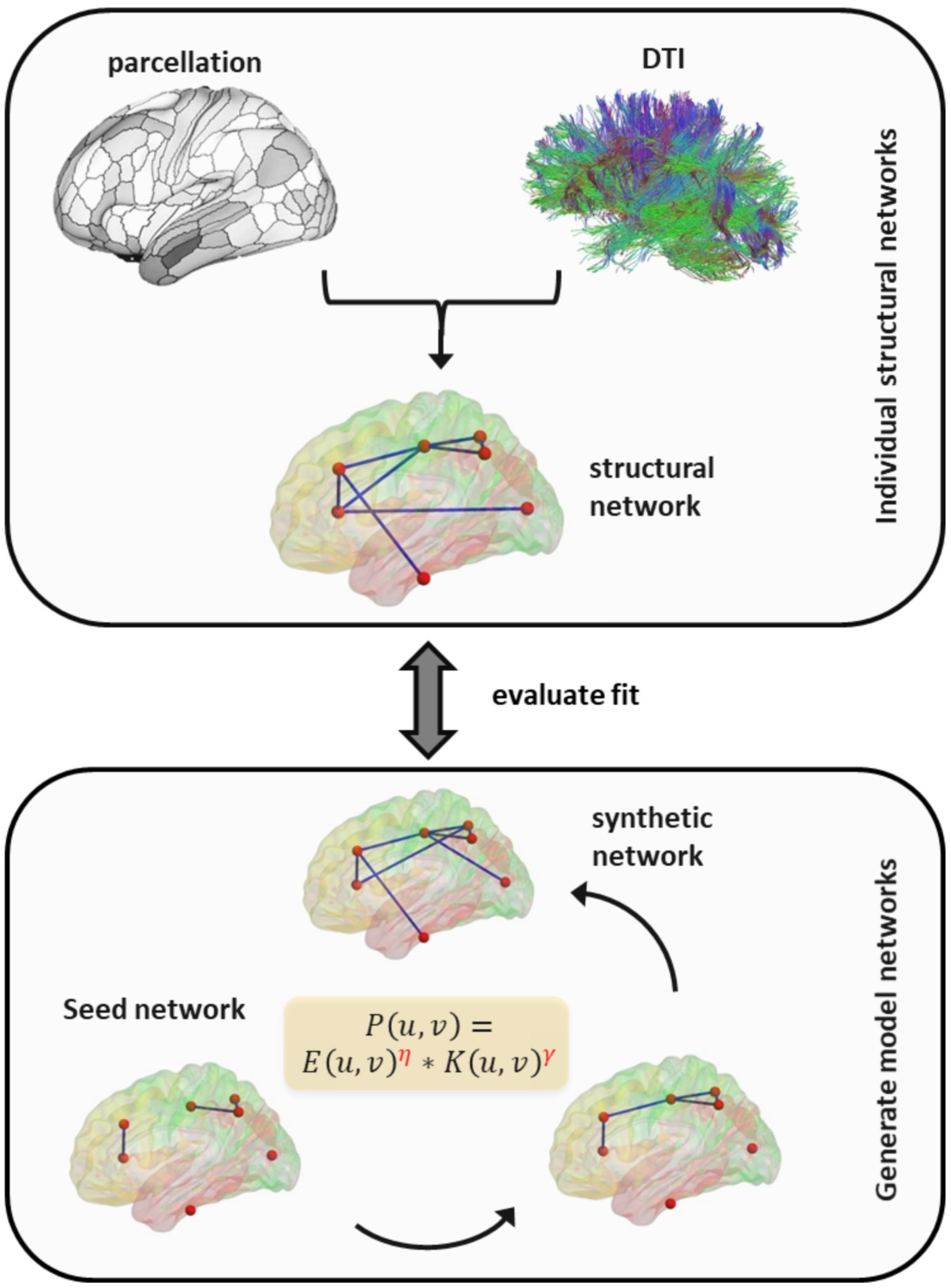
Overview of generative network models. Deterministic whole-brain fiber tracking was performed to reconstruct white mater pathways, from which we constructed structural networks linking 180 regions of interest. By retaining edges present in all subjects, a seed network was created and then edges were added stepwise with the probability of edge formation P(u,v) until the number of connections in the synthetic network was the same as that in the observed structural network. The fitness of the synthetic network was evaluated by comparing the degree, clustering coefficient, betweenness centrality and edge length distributions between the synthetic network and the observed structural network.

## METHODS AND MATERIALS

### Participants

We studied 152 healthy controls (HC) without a first-degree relative with mental illness (mean [SD] age, 30.32 [10.28] years; 94 women), 32 unaffected first-degree relatives (REL) of schizophrenia patients (33.25 [11.50] years, 19 women) and 66 unrelated patients satisfying DSM-IV-TR criteria (see Supplement) for schizophrenia (SZ, 32.77 [9.26] years; 20 women) in Mannheim, Germany. All participants provided written informed consent for the protocols approved by the local Ethics Committee of the University of Heidelberg.

### Neuroimaging data acquisition and processing

Diffusion Tensor Imaging (DTI) data were acquired with a 3-T Siemens Trio scanner using two echo planar imaging (EPI) sequences with different parameters (see Supplement for details). DTI data were preprocessed with standard routines implemented in the software package FSL (https://fsl.fmrib.ox.ac.uk/fsl/). The full pipeline included the following steps: correction for head motion and eddy currents by affine registration to *b*0 image, extraction of non-brain tissues (26), and linear diffusion tensor fitting. After estimating the diffusion tensor, we performed deterministic whole-brain fiber tracking using a modified FACT algorithm (27). Further methods details are provided in the Supplement.

### Construction of generative network models

We constructed synthetic networks using generative models. After defining a seed network consisting of those edges common across all subjects, edges were added one at a time until the number of edges in the synthetic network conformed to that of the observed network. The relative probability of edge formation was evaluated at each step according to the equation:

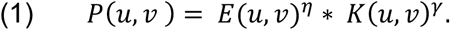

Here *E*(*u,v*) denotes the fiber distance between brain areas *u* and *v*, and *η* controls the edge length. When *η* is negative, short-distance edges are favored, while a positive *η* favors long-distance edges. The term *K*(*u,v*) represents the topological relationship between brain areas *u* and *v*, and *γ* represents its relative importance. Notably, *K*(*u,v*) can be varied to realize different wiring rules.

In this study, we limited our analysis to four generative models, each representing one of four previously-studied classes: the geometric model, the degree-product model, the clustering-product model and the matching index (MI) model (20). In the geometric model, the probability of connectivity *P*(*u,v*) between regions *u* and *v* is a function of the distance *E*(*u,v*) between them. In the degree-product and the clustering-product model, the connection probability function includes an additional topological term *K*(*u,v*), which is the product of degrees (number of connections of a brain region) or clustering coefficients (fraction of connection triangles around a brain region) between nodes *u* and *v* respectively. In the matching index model, *K*(*u,v*) denotes the normalized number of nearest neighbors in common between two nodes (homophily). The topological parameters were computed using the Brain Connectivity Toolbox (https://sites.google.com/site/bctnet/Home) as implemented in MATLAB.

To evaluate the fitness of synthetic networks and to optimize models, we define an energy function that measures how dissimilar a synthetic network is to the observed network:

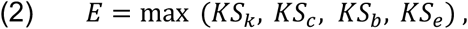

Here, each term is a Kolmogorov-Smirnov statistic that compares degree (*k*), clustering coefficient (*c*), betweenness centrality (*b*) and edge length (*e*) distributions of synthetic and observed networks. Since we defined energy as the maximum of the four statistics, smaller energy indicated greater fitness.

We used classical Monte Carlo methods to find the parameters (*η,γ*) that generated networks with minimal energy. The procedure starts from randomly sampling 2000 points from the defined parameter space. Then, by computing the energy at each point and dividing the whole parameter space into Voronoi cells, we sample points preferentially within Voronoi cells with low energy. We repeated this procedure five times until it converged to a (locally) optimal solution (20).

### Cognitive assessment and factor construction

In a subset of 120 individuals from all three groups, we assessed a range of cognitive subdomains frequently impaired in schizophrenia (including attention and psychomotor speed, executive function, memory, impulsivity and social emotional cognition) using the Cambridge Neuropsychological Test Automated Battery (CANTAB) (28, 29). Considering that the CANTAB measures were correlated with each other, we performed a principal component analysis (PCA) to reduce the redundancies and minimize potential for Type I error (30). The first component accounted for 27.1% of the variance and factor loadings were consistently negatively correlated with correct response rates and positively correlated with latency (or reaction times) across the seven test domains, suggesting that *lower* factor values indicate better individual cognitive performance. The detailed description of methods and a full list of included outcome measures across tasks and the resulting cognitive factors are provided in the Supplementary Information.

### Polygenic risk score

We used standard methods to extract genomic DNA from EDTA blood to perform genome-wide SNP (single nucleotide polymorphism) genotyping of all individuals using the Infinium PsychArray (Illumina Inc). Quality control and imputation was performed with Gimpute (31) (see Supplement for details). To control for population stratification, we performed a PCA on the linkage-disequilibrium pruned set of autosomal SNPs using GCTA (32). Then we excluded outliers and used the first five principal components as covariates in the following association analyses of model parameters. We computed the polygenic risk score with PRSice v-2, while the expected value of the missing genotypes were imputed based on the sample allele frequency (33). In this study, genome-wide association (34) nominal *P* < 0.05, was used to achieve a balance between the number of false-positive and true-positive risk alleles (35, 36). The association analyses were repeated for thresholds of nominal *P* < 0.01 and of nominal *P* < 0.1.

### Statistical analysis

For each participant, we tuned the parameters (*η, γ*) to the space where the generative model always produced synthetic networks with near-lowest energy. Within this space, we analyzed the top 1% minimal-energy synthetic networks. We compared the individual averaged energy of 1% lowest-energy synthetic networks between different types of generative models and different groups of participants using a one-way analysis of variance (ANOVA) with the Statistical Package for the Social Sciences (SPSS) 24. Then, we compared the parameters of the best-fitting model between groups using a general linear model. To investigate the genetic association of the model factors and to circumvent the potential effect of confounding factors not related to the genetic risk for the disorder in patient populations (37), we assessed the correlation between the parameters of the best-fitting model and polygenic risk scores in healthy controls only while controlling for age, sex, DTI protocol, temporal signal to noise ratio (tSNR) (38) and the first five principal components of population structure. To investigate the behavioral relevance of the identified model parameters, we assessed the correlation between the parameters of the best-fitting model and individual cognitive factor loadings while controlling for age, sex and tSNR in the three groups separately. To investigate whether our best-fitting model could capture other structural network abnormalities in schizophrenia, we also computed the global efficiency, modularity and hub degree (the degree of the top 10% highest-degree nodes) using the Brain Connectivity Toolbox. We also investigated the effect of antipsychotics on our model parameters (see Supplement).

## RESULTS

### Sample characterization

The groups were matched for age, education, tSNR and head motion, but not for sex and acquisition protocol (see detailed demographic and clinical characteristics as well as image quality control parameters in Table 1). To account for the group differences in these latter variables, we included sex and DTI protocol as covariates of no-interest in all analyses that included multiple groups.

**Table 1:** Demographic, clinical and neuroimaging characteristics

| | Healthy controls<br>(n = 152) | First-degree relatives<br>(n = 32) | Schizophrenia patients<br>(n = 66) | F or $\chi^2$ value | P value |
| --- | --- | --- | --- | --- | --- |
| **Demographic characteristics** |  |  |  |  |  |
| Age (years) | 30.32 ± 10.28 | 33.25 ± 11.50 | 32.77 ± 9.26 | 1.977 | 0.141 |
| Sex (male / female) | 58/94 | 13/19 | 46/20 | 18.95 | < 0.001 |
| Acquisition protocol (32/12 channel coil) | 76/76 | 21/11 | 66/0 | 50.71 | < 0.001 |
| Education (years) | 15.40 ± 1.60 | 15.19 ± 2.34 | 14.86 ± 2.17 | 1.881 | 0.155 |
| **Clinical characteristics** |  |  |  |  |  |
| PANSS positive | n.a | n.a. | 14.61 ± 7.54 | - | - |
| PANSS negative | n.a | n.a | 14.51 ± 8.27 | - | - |
| PANSS general | n.a | n.a | 31.70 ± 11.42 | - | - |
| PANSS total | n.a | n.a | 60.82 ± 23.51 | - | - |
| Duration of illness (years) | n.a. | n.a. | 10.44 ± 8.34 | - | - |
| Olanzapine equivalents (n= 52) | n.a. | n.a. | 15.04 ± 8.91 |  |  |
| **QC parameters** |  |  |  |  |  |
| DTI: mean relative root-mean-square displacement (mm) | 0.31 ± 0.10 | 0.33 ± 0.12 | 0.35 ± 0.18 | 1.844 | 0.160 |
| DTI: tSNR | 5.85 ± 0.29 | 5.76 ± 0.25 | 5.85 ± 0.26 | 1.491 | 0.227 |

PANSS = Positive and Negative Syndrome Scale (62). tSNR = temporal signal to noise ratio, QC = quality control

### Generative Network models

Comparison between the four network models revealed significant differences in mean energy (repeated measure ANOVA: *F*(3,453) = 2964.277, *p* < 0.001) in HC, with the matching index (MI) model showing the lowest energy level (see Figure 2A). Importantly, when including all diagnostic groups in the analysis, the group by model type interaction was not significant (repeated measure ANOVA with group as between-subjects factor, sex and DTI protocol as covariates of no-interest: *F*(6,735) = 0.870, *p* = 0.516), arguing for the same pattern across all groups. Hence, in our following investigation we focused on the analysis of the MI model, as it provided the best fit to the individual, experimentally-derived structural networks across diagnostic groups.

**Figure 2.**
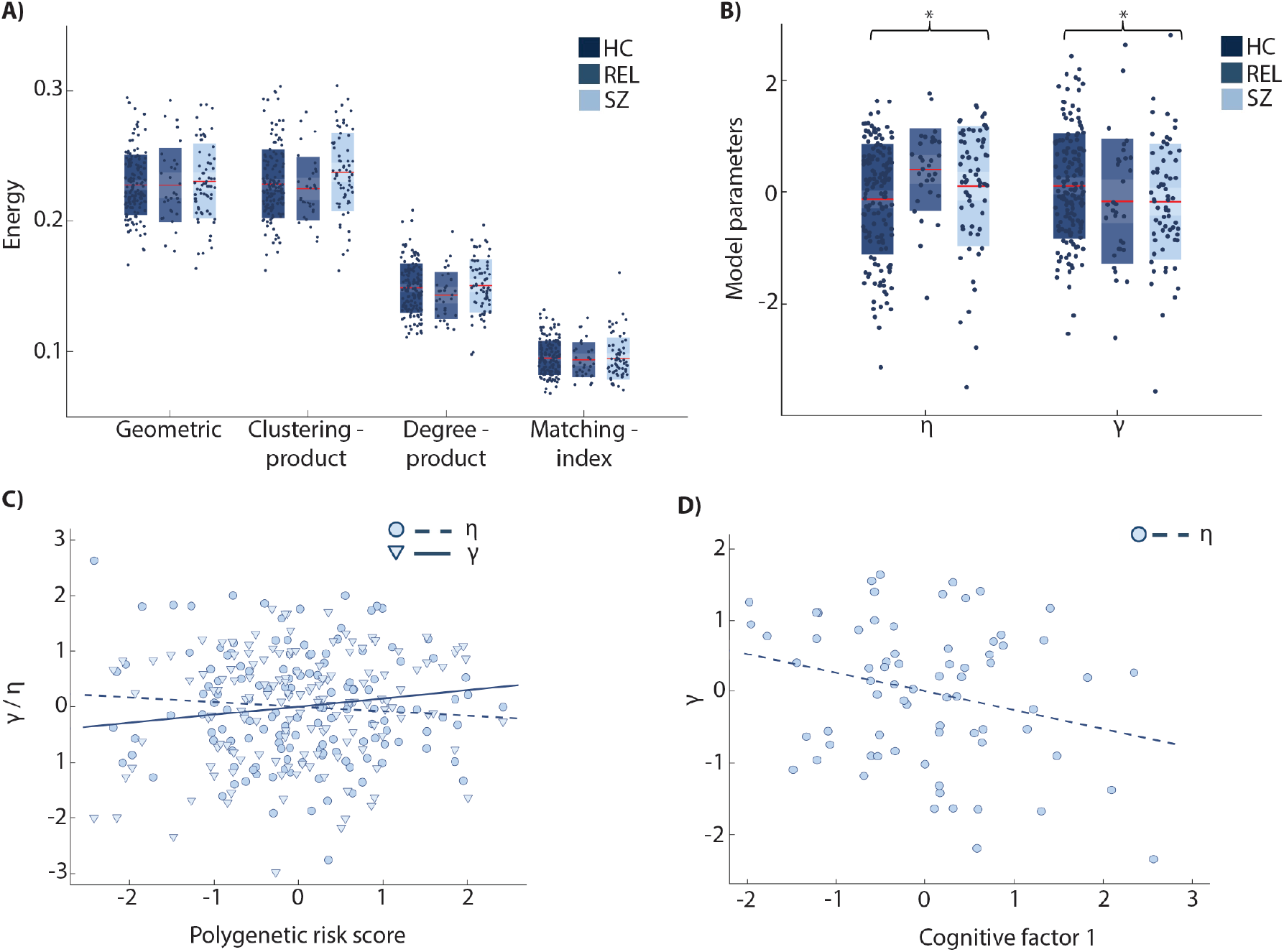
Generative model networks in health and disease. (A) Individual energy was significantly different among the four types of models (repeated-measure ANOVA: *F*(3,453) = 2964.277, *p* < 0.001) without a significant model by group interaction (*F*(6,735) = 0.870, *p* = 0.516). The matching index model showed the lowest energy. (B) In the matching index model, there was a significant between-group effect on the distance parameter η (ANOVA, *F*(2,245) = 4.777, *p* = 0.009) and on the topological parameter γ (ANOVA, *F*(2,245) = 3.054, *p* = 0.049) correcting for sex and DTI protocol. Red lines indicate mean values and boxes indicate one standard deviation of the mean. Asterisks denote significant difference between all diagnostic groups. (C) Individual polygenic risk score for schizophrenia was significantly positively associated with the distance parameter η (rpar = 0.173, p = 0.045) and trend-wise negatively associated with the topological parameter γ (rpar = −0.154, p = 0.073). (D) After performing principal component analysis on 13 main outcome measures of the neuropsychological test battery, we obtained five components whose eigenvalues were larger than 1, and we found a significant negative correlation between the first component and η (r = −0.261, p = 0.029) in healthy controls.

In Figure 3B, we illustrate the synthetic network structure of a single subject at different parameter values, thereby offering an intuition regarding the roles of each parameter in the MI model. As expected, the two parameters *η* and *γ* are strongly anti-correlated (Pearson correlation: *r* = −0.613, *p* < 0.001) in HC, a relation that was conserved across diagnostic groups (relatives: *r* = −0.535, *p* = 0.002, patients: *r* = −0.492, *p* < 0.001). Investigating the between-group difference of the two parameters, we found a significant between-group effect on the distance parameter *η* (ANOVA, sex and DTI protocol as covariates of no-interest; *F*(2,245) = 4.777, *p* = 0.009) and also on the topological parameter *γ* (ANOVA, same covariates of no-interest; *F*(2,245) = 3.054, *p* = 0.049, see Figure 2B). Post-hoc analysis confirmed significant differences between HC and SZ (η: *F*(1,214) = 3.956, *p* = 0.048; *γ*: *F*(1,214) = 4.707, *p* = 0.031) as well as between HC and REL in *η* (*F*(1,180) = 8.970, *p* = 0.003), but not in *γ* (*F*(1,180) = 2.609, *p* = 0.108). We found no significant correlation between individual olanzapine equivalents and model parameters (*η*: *r* = −0.121, *p* = 0.393; *γ*: *r* = 0.092, *p* = 0.517).

**Figure 3.**
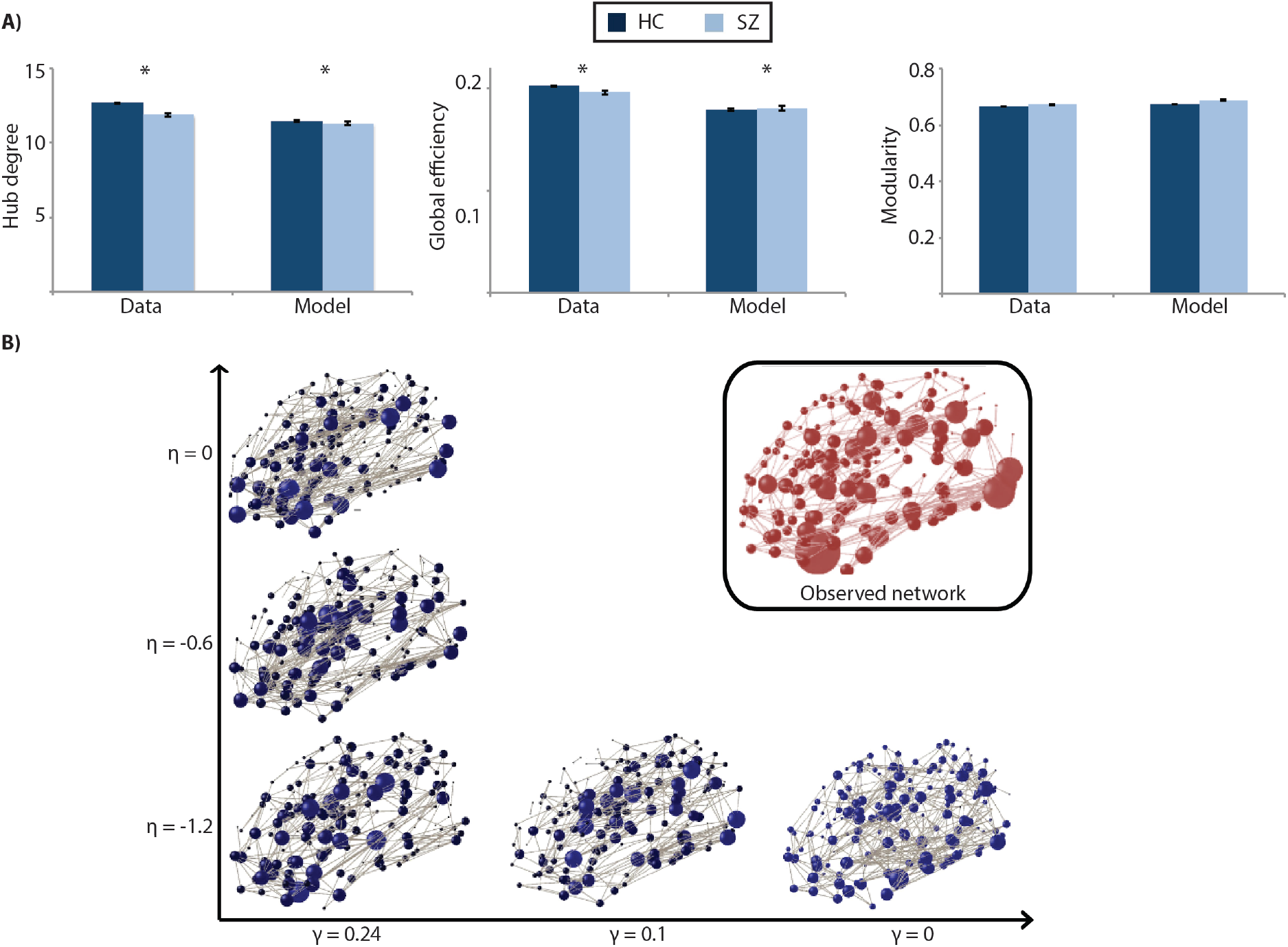
Topological characteristics in health and disease. (A) Comparison of several network characteristics in data and model between healthy controls (HC) and schizophrenia patients (SZ). The matching index model captured the abnormal hubness and global efficiency in schizophrenia patients well, while also adequately modeling no modularity difference between groups. The deep blue bars represent the clustering coefficient, modularity and hub degree of the synthetic network (model) and observed network (data) in healthy controls, respectively, while the light blue bars represent schizophrenia patients. Bars indicate mean values. Error bars indicate standard errors. Asterisks denote significant difference between diagnostic groups. Note that raw data is displayed. (B) Visualization of parameter effects on network structure for a single subject. The topological parameter γ mainly influences the degree distribution with larger γ corresponding to the occurrence of more and larger hubs, while the distance parameter η appears to tune the spatial position of hubs in the network. Here, η = −1.2 and γ = 0.24 correspond to the best-fitting model.

When comparing global efficiency, modularity and hub degree between HC and SZ, we detected significant between-group differences in global efficiency (ANOVA, same covariates of no-interest; *F*(1,214) = 11.143, *p* = 0.001) and in hub degree (ANOVA; *F*(1,214) = 8.602, *p* = 0.004) in the observed networks. In the synthetic networks, we also detected a trend-wise significant between-group difference in hub degree (ANOVA; *F*(1,214) = 3.218, *p* = 0.074), and a significant difference in global efficiency (ANOVA; *F*(1,214) = 8.082, *p* = 0.005).

No group differences were found in modularity for neither the observed data (ANOVA; *F*(1,214) = 0.100, *p* = 0.752) nor the synthetic network (ANOVA; *F*(1,214) = 0.251, *p* = 0.617, see Figure 3A).

### Polygenic risk score

To characterize the influence of genetic risk for schizophrenia on both model parameters, we correlated the individual participants, risk scores with *η* and *γ*. We found a significant positive association for the distance parameter *η* (*r*_par_ = 0.173, *p* = 0.045) and a weaker, trend-wise negative association for the topological parameter *γ* (*r*_par_ = −0.154, *p* = 0.073, Figure 2C) in HC for all genetic variants with a nominal genome-wide significant association to schizophrenia (*P* < 0.05, uncorrected). Supplemental analyses confirmed the robustness of this finding to the choice of significance threshold used for polygenic risk score computation: nominal *P* < 0.01: *η*(*r*_par_ = 0.154, *p* = 0.074), *γ* (*r*_par_ = −0.156, *p* = 0.071); *P* < 0.1: *η* (*r*_par_ = 0.196, *p* = 0.022), *γ* (*r*_par_ = −0.174, *p* = 0.043).

## CANTAB

Exploring the correlation between the individual participants, scores of the first component and the distance parameter *η* (or the topological parameter *γ*), we found a significantly negative association for *η* (*r* = −0.261, *p* = 0.029) and no association for *γ* (*r* = 0.108, *p* = 0.374) in HC. No association was found in relatives (*η*: *r* = −0.137, *p* = 0.589; *γ*: *r* = 0.035, *p* = 0.890) and in patients (*η*: *r* = 0.261, *p* = 0.240; *γ*: *r* = −0.133, *p* = 0.556, Figure 2D). As expected, healthy controls showed lower factor loadings compared to relatives and patients (*F*(2, 115) = 7.680, *p* = 0.001). We did not detect any correlation between the other four cognitive components and the network model parameters.

## DISCUSSION

Converging evidence points towards alterations in the structural connectome of schizophrenia patients, but the mechanistic disturbances in the process of network formation leading to those remain poorly understood. Here we replicate prior work on the performance of generative network models (20) and thereby demonstrate that this approach yields synthetic networks that simulate many properties of structural brain networks in health and disease. We further identify significant differences between healthy controls, first-degree relatives and schizophrenia patients for the two best-fitting model parameters in a pattern that aligns with the increasing risk for schizophrenia. Specifically, these differences imply lesser geometrical constraints and lesser complexity of topological facilitation of network formation in schizophrenia. Moreover, by demonstrating differential associations between polygenetic risk score and the network formation parameters, we provide a potential mechanistic explanation of how genetic risk contributes to the malformation of brain networks. These parameters also demonstrated behavioral relevance by predicting latent features of cognitive functioning.

Firstly, our data replicate previous accounts of a superior performance of a model – parameterized by spatial distance and a matching index – in capturing several topological features of structural brain networks in comparison to other models (20). This observation aligns well with the current theory that brain networks are shaped by a mixture of geometrical constraints and topological complexity. The human brain is physically expensive and the metabolic cost of building and maintaining the axonal connections increases with connection length (39). Therefore, cost minimization seems to be an important evolutionary rule for many aspects of brain anatomy (40–42). However, cost minimization alone only poorly accounts for some of the adaptive properties of the human connectome, such as the capacity for information processing (43), which should therefore result from a trade-off between wiring cost and formation of topological features (11). In our best-fitting model, this trade-off is represented by the two generative factors: the distance parameter *η* penalizes the formation of long connections by imposing a spatial constraint, while the topological parameter *γ* favors connections between regions sharing similar neighbors, meaning that cortical areas that have similar inputs and outputs tend to be connected, compatible with Hebb’s law. Interestingly and in contrast to a previous study (20), we observe a negative correlation between our model parameters *η* and *γ*, suggesting that connectomes that are less constrained by spatial distance also exhibit a lower topological neighborhood structure. Together, our findings thus rather argue against a simple trade-off between both parameters. Instead our data may indicate that connectomes may be better modeled with a single, to-be-discovered latent factor that could account for both parameters simultaneously.

Secondly, when extending this framework to model mechanisms of connectome formation in schizophrenia patients and first-degree relatives, we detected no significant between-group differences in the fit between the synthetic networks and the observed networks across models. Importantly, these results suggest that the same wiring rules can equally well describe both normal and abnormal brain network formation. Previous studies have found increased connection distance of brain networks (12, 13) and changed network topology including reduced clustering and modularity (14, 15, 44) in schizophrenia (see Figure 3A). In line with this, theoretical accounts suggest that the abnormal organization of brain networks in schizophrenia may result from a biased trade-off between generative factors of homophilic attraction and distance penalization in the process of brain network formation (11). To probe further, we tested whether individual model parameters contributed differently to network formation in all three study groups. We identify smaller values of the distance parameter *η* in HC than in relatives and patients, while values of the topological parameter *γ* were higher in HC than in relatives and patients. This said, larger *η* values in relatives and patients indicate a lower distance penalization, thus increasing the edge length distribution, which is consistent with the increased connection distance of brain networks in schizophrenia (12, 13). Since the topological parameter mainly influences the degree distribution (21) (see Figure 3B), smaller *γ* values in patients and relatives indicate the presence of fewer and smaller hubs. This corroborates previous findings that suggested brain networks in schizophrenia are less clustered and have fewer hubs (9, 13). Notably, the presence of decreased spatial constraints and homophilic association in a sample of unaffected first-degree relatives suggests that these network mechanisms may resemble intermediate phenotypes, i.e., brain phenotypes that relate to the increased familial risk for schizophrenia rather than being epiphenomena of potential confounds such antipsychotic medication.

Thirdly, we examined the associations between model parameters and schizophrenia polygenic risk in healthy individuals. We identified a positive correlation between polygenic risk and the distance parameter *η*. In addition, we showed a trend-wise negative association between polygenic risk and the topological parameter *γ*, suggesting that increasing genetic risk load for schizophrenia leads to a diminished distance penalization and local homophily of the structural brain connectome. In general, two interconnected factors are thought to contribute to the formation of long distance connections in the brain: a) neurons connect to each other at an early stage of neurodevelopment when the neural system is small in scale, and b) at later stages, axons follow developmental pathways already established by earlier pioneer neurons (fasciculation) (45). Studies in *C. elegans* show that 70% of long distance connections are already present at the time of hatching (46). This observation suggests that most neurons already form early connections in a spatially localized system where guidance through differential expression of guidance molecules, such as *netrin* (47) and *slit* (48), is feasible. In contrast, these molecules will not be the preferred driver to guide axons at later stages of development because source and projection neurons are further away (45). Importantly, the genes coding for such guidance molecules have been repeatedly implicated in the pathophysiology of schizophrenia (49–51). It is interesting to speculate that altered spatial expression of guidance molecules imposes less spatial constraints on brain network formation, and these fine-scale mechanisms are reflected in our large-scale model parameters.

Moreover, we found a negative correlation between the distance parameter *η* and individual scores of the first principal cognitive factor, predominantly capturing converging aspects of executive function and memory, in HC. In particular, we observed that larger values of *η* (reflecting less geometric constraints, thus a higher probability of long distance connections) were associated with better cognitive performance. Healthy human brain networks usually contain only a small fraction of long-distance shortcuts preferentially linking hub regions (12, 52). Although these long-distance connections are expensive in terms of material and metabolic cost, they greatly reduce the path-length of information transfer between spatially remote regions, thus increasing the potential for efficient information processing (11, 53), but not in weighted networks with weights based on streamline density (54). A number of previous studies have shown that more topologically efficient structural and functional networks are associated with increased IQ (55, 56) and cognitive performance (57, 58). Therefore, while long-range connections are beneficial for brain function (59), they may also be subject to increased vulnerability to disease (60). Indeed, previous studies have shown an increased proportion of long-range connections in schizophrenia resulting in brain networks shifted towards random networks (61). The association of the distance parameter *η* and cognition was not detectable in schizophrenia patients and first-degree relatives, suggesting an optimum in the number of long-range connections potentially exceeded in those populations.

There are a number of methodological considerations that deserve discussion. Firstly, since our goal was to investigate the mechanisms underlying the abnormal brain network formation in schizophrenia with GNM, we restricted our analysis to an already validated model framework (20). In principle, other wiring rules may be used to model connectome formation and evaluate the fitness of the resulting models. However, such an exploration is beyond the scope of this paper and would require extensive prior validation in healthy control datasets. Secondly, even complete correspondence of two networks does not necessarily imply that both models have been shaped by the same biological mechanism(s). While we have attempted to limit interpretational restraints by external validation with other well-established network features, it is important to note that generative models can be used to offer candidate mechanisms for an observed topology, but cannot conclusively prove that a given candidate mechanism actually occurred in the developing organism (16). Thirdly, while GNMs can provide insights into the formation of structural brain networks, they do not explicitly model neurodevelopmental processes. Such investigations are warranted, and will require longitudinal datasets as well as the use of an advanced GNM framework explicitly modeling variant developmental processes within subjects.

In conclusion, we show that the distinct wiring rules can simulate normal and abnormal network formation in humans, identify intermediate connectomic phenotypes for schizophrenia familial risk manifesting as altered spatial and topological characteristics of brain connectome formation in schizophrenia and first-degree relatives. We further corroborate the link of these network mechanisms to schizophrenia polygenic risk and demonstrate their relevance for individual cognitive function in a domain frequently disturbed in schizophrenia. Together, these data suggest that brain network formation is under strong genetic control, is optimized to support cognitive functioning and disturbed in heritable developmental disorders such as schizophrenia. While these results provide important insight into the wiring mechanisms in health and schizophrenia, longitudinal studies in developmental cohorts are needed to further elucidate successful and aberrant brain connectome formation.

## ACKNOWLEDGEMENTS

The authors thank all individuals who have supported our work by participating in our studies. There was no involvement by the funding bodies at any stage of the study. We thank Ilka Alexi, Oliver Grimm, Leila Haddad, Maria Zangl, Franziska Danneberg and Mathias Kienow for valuable research assistance.

X.Z. is a Ph.D. scholarship awardee of the China Scholarship Council. U.B. acknowledges grant support by the German Research Foundation (DFG, BR 5951/1-1). H.T. acknowledges support from the German Federal Ministry of Education and Research (BMBF, grant 01GQ1102) and the German Research Foundation (DFG, GRK 2350 project B2; TO 539/3-1). DH acknowledges grant support by the German Research Foundation (DFG HI 1928/2-1). AML acknowledges grant support by German Federal Ministry of Education and Research (BMBF, grants 01ZX1314G, 01GS08147, 01GQ1003B, Collaborative Research Center 1158 subproject B09), European Union’s Seventh Framework Programme (FP7, grants 602450, 602805, 115300 and HEALTH-F2-2010-241909, Innovative Medicines Initiative Joint Undertaking (IMI, grant 115008), German Research Foundation (DFG, ME 1591/4-1) and Ministry of Science, Research and the Arts of the State of Baden-Wuerttemberg, Germany (MWK, grant 42-04HV.MED(16)/16/1). DSB and RFB would like to acknowledge support from the John D. and Catherine T. MacArthur Foundation, the Alfred P. Sloan Foundation, the Army Research Laboratory and the Army Research Office through contract numbers W911NF-10-2-0022 and W911NF-14-1-0679, the National Institute of Health (2-R01-DC-009209-11, 1R01HD086888-01, R01-MH107235, R01-MH107703, and R21-M MH-106799), the Office of Naval Research, and the National Science Foundation (BCS-1441502, CAREER PHY-1554488, and BCS-1631550). The content is solely the responsibility of the authors and does not necessarily represent the official views of any of the funding agencies. ES gratefully acknowledges grant support by the Deutsche Forschungsgemeinschaft, DFG (SCHW 1768/1-1). R.F.B. was supported by Indiana University Office of the Vice President for Research Emerging Area of Research Initiative, Learning: Brains, Machines and Children. MMN acknowledges grant support by the German Federal Ministry of Education and Research (BMBF) through the Integrated Network IntegraMent (Integrated Understanding of Causes and Mechanisms in Mental Disorders), under the auspices of the e:Med Programme (grant 01ZX1314A/01ZX1614A to M.M.N.).

