## Supplemental Material for "Generative network models identify biological mechanisms of altered structural brain connectivity in schizophrenia"

***Supplementary Information***

### SUPPLEMENTARY METHODS

#### Participants

Patients were consecutively recruited from the Department of Psychiatry and Psychotherapy at the Central Institute of Mental Health in Mannheim, Germany. Diagnoses were made by staff psychiatrists. Clinical evaluation included ascertainment of personal and family history and detailed physical and neurological examination. Patients were excluded if: (i) they were aged <18 or >65 years, (ii) they had a history of brain trauma or neurological disease.. The local ethics committee (Medical Faculty at Heidelberg University, Germany) approved the study.

#### Neuroimaging data acquisition

Diffusion Tensor Imaging (DTI) data were acquired with a 3-T Siemens Trio scanner using two echo planar imaging (EPI) sequences with different parameters: 1) 32 channel multi-array head-coil, TE/TR = 86/8400 ms, 2 mm slice thickness, field of view (FOV) = 256*256 $mm^2$, 64 slices, and 46 diffusion directions at *b*-value of 1000 $s/mm^2$; 2) 12 channel coil, TE/TR = 86/14000 ms, 2 mm slice thickness, FOV = 256*256 $mm^2$, 64 slices, and 60 diffusion directions at *b*-value of 1000 $s/mm^2$. A total of 163 participants were scanned with the first sequence and 87 participants were scanned with the second sequence.

#### Deterministic fiber tracking

When performing deterministic whole-brain fiber tracking , we initiated 1,000,000 streamlines for each subject and removed those with a length of less than 10 mm. To construct the structural connectome, the cerebral cortex was parcellated into 360 areas (2) and the number of streamlines connecting every pair of brain areas was used as an estimate of structural connectivity. Note that if the number of streamlines connecting two regions was less than 5, we set the connection weight to zero to minimize bias due to false positives. In our analysis, we only focused on the right hemisphere (180 areas) following the precedent set in related prior work (3, 4). To compare the structural connectome between groups, every connection was required to be present in at least 70% of subjects (5).

**Principal component analysis of CANTAB measures**

In a subset of 120 individuals (74 healthy controls, 21 relatives and 25 patients), we assessed cognitive function using the Cambridge Neuropsychological Test Automated Battery (CANTAB) implemented on a touchpad tablet computer. The selected tests covered a range of cognitive subdomains frequently impaired in schizophrenia (6, 7) including attention and psychomotor speed, executive function, memory, impulsivity and social emotional cognition (Emotion Recognition Task (ERT), Pattern Recognition Memory (PRM), Spatial Span (SSP), Stocking of Cambridge (SOC), Reaction Time (RTI), Attention Switching Task (AST) and Information Sampling Task (IST)). We performed a principal component analysis (PCA) on the acquired measures to reduce the redundancies and minimize potential for Type I error in multiple comparisons (8). For this, we selected two main outcome measures per test (with the exception of SOC, for which only one outcome measure is available) and performed a PCA, which resulted in five components whose eigenvalues were larger than 1. The first component accounted for 27.1% of the variance and factor loadings captured mainly executive function and memory. The first component was consistently negatively correlated with correct response rates and positively correlated with latency (or reaction times) across the seven test domains, suggesting that *lower* factor values indicate better individual cognitive performance. The detailed description and a full list of included outcome measures across tasks and the resulting cognitive factors are provided in Table S1.

### Genotyping processing

Quality control (QC) and imputation was performed with Gimpute (9) including the following steps: Removal of SNPs with sex chromosome heterozygosity, a missing rate greater than 0.05, deviation from Hardy-Weinberg equilibrium in controls ($P$ < $10^{-6}$) and autosomal heterozygosity deviation of greater than 0.2 as well as removal of samples with a missing rate greater than 0.02. Phasing and imputation was conducted using SHAPEIT and IMPUTE2 (10-12) with the imputation reference panel from the 1000 Genome Project dataset (August 2012, 30,069,288 variants, release “v3.macGT1”). After imputation, we only retained SNPs with an imputation INFO score larger than 0.6, minor allele frequencies larger than 0.01 and successfully imputated in at least 20 individuals. The proportion of alleles shared identity-by-decent estimated using PLINK(13) (www.cog-genomics.org/plink/1.9/) was used to identify relatedness for all pairs of samples. A threshold of $\hat{\pi}$ > 0.2 was used to identify related pairs of samples and exclude one member of each pair at random.

### Olanzapine equivalents

To investigate the effect of antipsychotics on the results, we converted the daily doses of patients' antipsychotic medication to olanzapine equivalents (OLZe) according to the classical mean dose method presented by Leucht and colleagues (14). This method is based on the analyses of 13 oral second-generation antipsychotics, haloperidol, and chlorpromazine compared with olanzapine 1 mg/d. To obtain OLZe, we weighted the mean dose of each antipsychotic by the study's sample size and finally divided by the weighted mean olanzapine dose.

Table S1. Neuropsychological measures and respective factor loadings

| Measures | Factor 1 | Factor 2 | Factor 3 | Factor 4 | Factor 5 |
|---|---|---|---|---|---|
| SSP span length | -0.579 | | | -0.323 | 0.350 |
| SSP mean time | 0.371 | 0.329 | | -0.629 | |
| IST correct win | | 0.644 | | | -0.406 |
| IST error | | -0.507 | -0.340 | | 0.458 |
| PRM mean latency | 0.670 | | -0.332 | | |
| PRM correct | -0.491 | | 0.529 | | 0.345 |
| SOC problems solved | -0.338 | 0.620 | | | |
| RTI movement time | 0.379 | | 0.618 | | 0.360 |
| RTI reaction time | 0.714 | | 0.409 | | |
| AST correct trials | -0.442 | 0.313 | | 0.582 | 0.319 |
| AST mean latency | 0.729 | | | | |
| ERT mean latency | 0.605 | 0.468 | | | |
| ERT correct | -0.652 | | | | |

*Note:* ERT, Emotion Recognition Task; PRM, Pattern Recognition Memory; SSP, Spatial Span; SOC, Stocking of Cambridge; RTI, Reaction Time; AST, Attention Switching Task; IST, Information Sampling Task.
